# FastDedup A fast and memory-efficient tool for read deduplication

**DOI:** 10.64898/2026.04.29.721745

**Authors:** Raphaël Ribes, Céline Mandier, Alice Baniel

**Affiliations:** ISDM, Univ Montpellier, CIRAD, INSERM, Montpellier, France; ISEM, Univ Montpellier, CNRS, IRD, Montpellier, France

## Abstract

PCR duplicate removal is a critical first step in high-throughput sequencing pipelines, yet existing tools struggle with speed, memory, or correctness at modern dataset scales. We present `FastDedup`, a Rust-based FASTX deduplicator that transforms each read or read pair to a compact `xxh3` hash fingerprint, drastically reducing memory usage and binding most of the execution time to disk I/ O. Benchmarked against six competing tools on synthetic human WGS datasets up to 300 million reads, `FastDedup` consistently leads on paired-end data, running more than 10 times faster than `fastp`. It also outperforms all tools on uncompressed single-end data, deduplicating a million reads in a second. We additionally report correctness failures in `prinseq++` and `clumpify`. `FastDedup` is available under the MIT License via GitHub, Bioconda, and Cargo.

## 1 Context

With the rise of Next-Generation Sequencing (NGS) technologies, the volume of sequencing data has increased exponentially. With this surge in data, the development of new tools for processing and analyzing sequencing reads became necessary. One of the first and most critical steps in the analysis pipeline is read deduplication, which involves identifying and removing duplicate sequences that can arise from PCR amplification or sequencing errors. This is crucial when working with DNA-seq data, but it can introduce bias in RNA-seq data since the duplication could be biological in nature (e.g., highly expressed genes).

**Solid Black Line:** Represents a single sequenced DNA fragment (height reflects length).

**Red Arrow (R1):** Forward read (Read 1).

**Blue Arrow (R2):** Reverse read (Read 2).

**A) Not a duplicate:** Shows two distinct fragments where the start position of Read 1 is shared, but different fragment lengths lead to unique mapping positions for Read 2.

**B) PCR Duplicate:** Shows two exact fragments where the start position of Read 1 and Read 2 is shared.

Given a fragment *F*, we define the 5′->3′ sequence as *R*1 with a length of *r*_1_, and the 3′->5′ sequence as *R*2 with a length of *r*_2_. *R*1 is the prefix of *F*, the reverse complement of *R*2 is the suffix of *F*, and *r*_1_ = *r*_2_.

Therefore, a PCR duplicate occurs when *F*_1_ and *F*_2_ share the same starting and ending positions in the genome, and thus have the same length. In that case, *R*1_1_ and *R*1_2_ are identical, and *R*2_1_ and *R*2_2_ are identical.

This step is crucial because removing these artifacts prevents false positives during variant calling, as PCR polymerases can introduce errors during amplification cycles. If an error occurs early, it is propagated into multiple daughter molecules. When sequenced, these identical, error-carrying molecules will appear as multiple independent reads supporting a mutation that does not actually exist in the biological sample [1].

Deduplication also helps correct allele frequency bias, as not all DNA molecules amplify with equal efficiency due to factors such as GC content or fragment length. This uneven amplification can artificially inflate the coverage of certain alleles. Deduplication also improves *de novo* assembly, especially in whole-genome sequencing. When paired-end reads are mapped to estimate the order and intervening distance between contigs, duplicates can create false-positive connections between contigs or introduce conflicting connections [2].

Before presenting our tool, we will first review existing tools for read deduplication and their limitations.

## 2 Existing tools and limitations

### 2.1: Fastp

`fastp` is a widely adopted, all-in-one FASTQ preprocessor written in C++ that performs quality control, adapter trimming, and deduplication within a single pass [3,4]. Unlike alignment-based tools that require computationally expensive mapping and coordinate sorting, `fastp` operates directly on raw sequences.

To achieve high performance, `fastp` implements a probabilistic deduplication mechanism based on Bloom filters and multiple hash arrays. For each read, or read pair, the sequence is mapped to multiple integer hash values. If all corresponding positions in the hash arrays are already marked as positive, the read is flagged as a duplicate. This streaming, single-pass architecture avoids reading and writing intermediate files to disk, drastically reducing input/output (I/O) bottlenecks and resulting in ultra-fast execution speeds.

A significant advantage of `fastp` is its highly configurable memory management. Users can adjust the `--dup_calc_accuracy` parameter to scale the hash buffer size from 1 GB up to 24 GB, allowing the tool to run on both standard personal computers and High-performance Computing (HPC) clusters. This advantage is also its incovenient because the user would have to set up specific parameters for each dataset, which can be a problem for non-expert users and require trial and error to find the optimal settings. Because of this, we will use the default settings.

However, this probabilistic approach involves a minor trade-off in absolute precision, as `fastp` has a theo-retical false-positive rate. Furthermore, `fastp`’s deduplication algorithm relies heavily on the exact matching of sequence lengths and coordinate regions. Consequently, applying certain quality trimming operations (such as sliding-window trimming via `--cut_front` or `--cut_tail`) prior to deduplication can interfere with the duplicate identification process, as trimmed reads may no longer perfectly match their duplicate counterparts.

We will be using the 1.1.0 version in this study.

### 2.2: FastUniq

`FastUniq` [5] is a deterministic deduplication tool explicitly designed for paired-end (PE) short reads. By directly comparing the nucleotide sequences of read pairs, it provides an exact identification of PCR duplicates.

Algorithmically, `FastUniq` operates by loading the entirety of the paired reads into memory, sorting them based on their sequences using a merge sort algorithm, and identifying duplicates by sequentially comparing adjacent read pairs in the sorted list.

A notable feature of its comparison logic is its ability to handle reads of varying lengths: a shorter sequence is considered identical to a longer one if it perfectly matches its 5′ prefix. If duplicates are found, the pair containing the longest reads is retained.

While `FastUniq` guarantees absolute precision for exact duplicate removal, its architectural design introduces critical resource bottlenecks. Because it relies on importing the entire dataset into RAM simultaneously using a hierarchical storage structure, its memory footprint scales linearly with the input file size. Benchmarking studies have highlighted this as a severe limitation for modern high-throughput data. For instance, processing a moderately sized dataset of approximately 69 million read pairs required over 50 GB of RAM [6].

While highly accurate, `FastUniq` is often impractical for massive datasets without access to HPC clusters. Furthermore, its scope is strictly limited to paired-end data, and it does not tolerate sequence mismatches.

We will be using the 1.1 version in this study.

### 2.3: Clumpify

`clumpify`, a tool from the Java-based BBTools suite [7], introduces an algorithmic paradigm to duplicate removal based on *k*-mer grouping. Rather than relying on exact sequence hashing or global coordinate sorting, `clumpify` identifies reads that share common *k*-mers and groups them into localized clusters or “clumps”.

This spatial reorganization provides a unique secondary benefit. By physically placing highly similar sequences adjacent to one another in the output file, `clumpify` dramatically improves standard `gzip` compression efficiency, as the compression algorithm can utilize much closer reference pointers.

A major functional advantage of `clumpify` is its high sensitivity to optical duplicates. By parsing the flowcell tile and pixel coordinates embedded within Illumina FASTQ headers, `clumpify` (via parameters such as `dupedist`) can explicitly distinguish between amplification-derived PCR duplicates and sensor-derived optical duplicates. For paired-end data, `clumpify` predominantly uses the first read (R1) to determine the clump, allowing the paired read (R2) to passively follow its counterpart, which maintains file synchronization while minimizing computational overhead.

To manage memory, `clumpify` implements a multi-phase strategy (KmerSplit and KmerSort). If the dataset exceeds available RAM, the tool can divide the data into temporary groups on the disk, sorting them individually before merging.

However, its reliance on the Java Virtual Machine (JVM) can lead to instability and high memory overhead on massive datasets. By default, high-memory tools within the BBTools suite will attempt to allocate all available system RAM [7], and JVM garbage collection overhead can cause severe bottlenecks unless strict multi-threading and heap-size limits are manually tuned.

We will be using the 39.79 bbmap version in this study.

### 2.4: SeqKit

`SeqKit` [8,9] is a comprehensive, cross-platform, and ultra-fast toolkit for FASTA and FASTQ file manipulation. Written in Go, it benefits from efficient concurrency management and the ability to produce static binaries with no external dependencies, making it highly accessible and stable across different operating systems.

For PCR deduplication, `SeqKit` provides the `rmdup` subcommand. This tool relies on an exact sequence matching approach using a hashing strategy. To process massive datasets without exhausting system memory, `seqkit rmdup` offers an MD5 mode (invoked via the `-m` or `--md5` flag). In this mode, the algorithm computes and stores only the MD5 digest of each sequence in the RAM rather than the full nucleotide sequence. This extreme memory efficiency allows `SeqKit` to maintain a very low resource footprint, even outperforming newer tools (e.g `prinseq++`) when handling vast amounts of single-end data.

Despite its speed and memory efficiency, `SeqKit rmdup` has a notable architectural limitation: it does not natively support the synchronized deduplication of paired-end reads. The `rmdup` command evaluates and processes individual files independently. It checks for duplicates inside each file separately; therefore, if *R*1_1_ = *R*1_2_ but *R*2_1_ ≠ *R*2_2_, *R*1_2_ will be flagged as a duplicate and removed while *R*2_2_ is retained, leading to data loss and desynchronization of the paired files. While workarounds exist using other `SeqKit` subcommands (such as `seqkit pair` to resynchronize files), `seqkit rmdup` is generally considered the optimal choice strictly for single-end libraries. This is why we will not use it for paired-end benchmarking in this study.

We will be using the 2.13.0 version in this study.

### 2.5: CD-HIT-DUP

`CD-HIT-DUP`, a specialized tool within the broader CD-HIT package [10,11], is designed to identify and remove both identical and nearly-identical duplicates. Unlike tools restricted to exact string matching, `CD-HIT-DUP` is particularly robust against sequencing errors. It incorporates a greedy incremental clustering algorithm that tolerates a defined threshold of mismatches, including both substitutions and insertions/deletions. `CD-HIT-DUP` relies on a prefix-suffix comparison approach. The tool first bins reads that share an identical prefix of *k* nucleotides. To accelerate this binning and sorting process, it uses a dual numeric encoding strategy, allowing prefixes of up to 27 nucleotides to be compactly represented as 64-bit integers. Within each bin, sequences are sorted by decreasing length, with the longest sequence acting as the cluster’s seed. The algorithm then evaluates the suffixes of the remaining sequences against this seed, utilizing a *k*-mer word-based index table to statistically predict sequence identity and bypass computationally expensive full pairwise alignments when possible. For paired-end libraries, `CD-HIT-DUP` conceptually joins the read pairs into a single sequence and checks prefixes at both ends simultaneously to ensure paired-end integrity.

We will be using the 4.8.1 cd-hit-auxtools version in this study.

### 2.6 PRINSEQ++

`prinseq++` [12] is an optimized C++ implementation of the Perl-based `prinseq-lite` program. It is designed to perform a comprehensive suite of quality control, filtering, trimming, and reformatting tasks for genomic and metagenomic sequence data.

For sequence deduplication, `prinseq++` departs from exact string comparison or sorting heuristics by implementing a probabilistic data structure known as a Bloom filter. During execution, each sequence is transformed using multiple fast, non-cryptographic hash functions, and the corresponding indices in a shared bit-array are flagged. If a newly evaluated sequence maps entirely to bits that are already set to positive, it is classified as a duplicate. This probabilistic model is highly advantageous for multi-threaded environments because it enables asynchronous read and write access to the filter with minimal thread blocking. Additionally, `prinseq++` natively supports the direct reading and writing of `gzip`-compressed FASTQ and FASTA files without the need for intermediate disk I/O, which significantly reduces hard drive storage requirements.

It also natively supports paired-end reads by processing pairs simultaneously and outputting synchronized R1 and R2 files. However, the deduplication function (`-derep`) is strictly limited to exact duplicates and does not tolerate sequencing errors. Furthermore, the probabilistic nature of the Bloom filter introduces a theoretically non-zero rate of false positives. Finally, when `prinseq++` is executed with multiple threads, the output sequences are not guaranteed to remain in the same order as the input files, which may require consideration if downstream tools expect strict global index synchronization.

We will be using the 1.2.4 version in this study.

### 2.7: SAM/BAM ecosystem tools

In addition to tools that operate directly on raw FASTQ sequences, the bioinformatics ecosystem relies heavily on utilities designed for SAM and BAM formats. These include PCR deduplication tools such as `samtools` (via `markdup`) [13] and Picard’s `MarkDuplicates` [14].

While these tools are typically used downstream after mapping reads to a reference genome, it is technically possible to convert raw, unaligned FASTQ files directly into unmapped SAM/BAM files to process them through these pipelines. However, even without the computationally heavy step of genomic alignment, the sheer format conversion from FASTQ to SAM/BAM introduces significant and unnecessary I/O and processing overhead. Because our primary objective is to evaluate and develop a highly optimized, memory-efficient tool strictly for direct FASTQ-to-FASTQ deduplication, these format-dependent tools fall outside the scope of our comparative analysis.

## 3 FastDedup: High-Performance FASTX Deduplication

The goal behind the development of `FastDedup` (or `FDedup`; Figure 2 Figure 3) was to create an exact FASTX PCR deduplication tool that prioritizes maximum speed and memory efficiency. To achieve this, the tool relies on `xxh3` [15,16], a non-cryptographic hash function, to compute a unique fingerprint for each read. These fingerprints are cached in memory using `fxhash` [17], which provides a low memory footprint.

**Figure 1:**
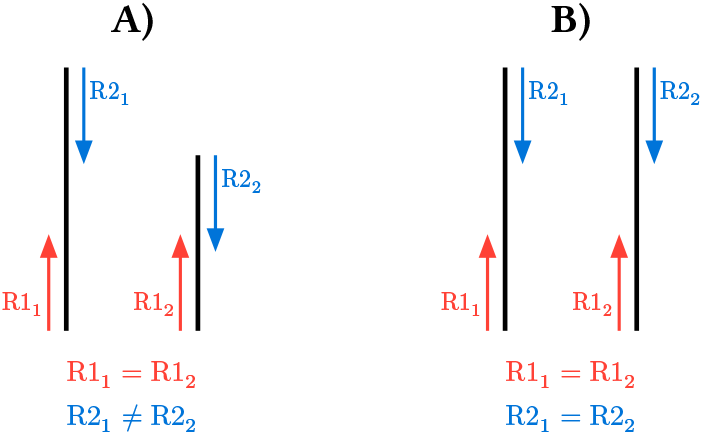
Identifying PCR Duplicates in Paired-End Sequencing.

**Figure 2:**
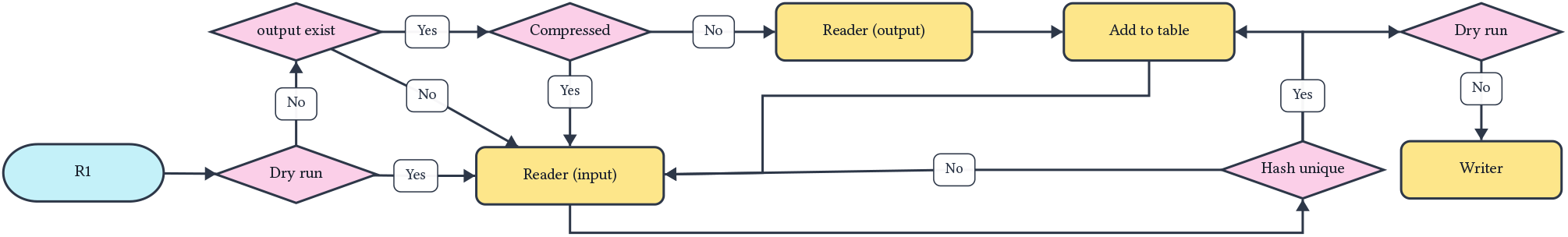
Architectural design of `FastDedup` for single-end reads.

**Figure 3:**
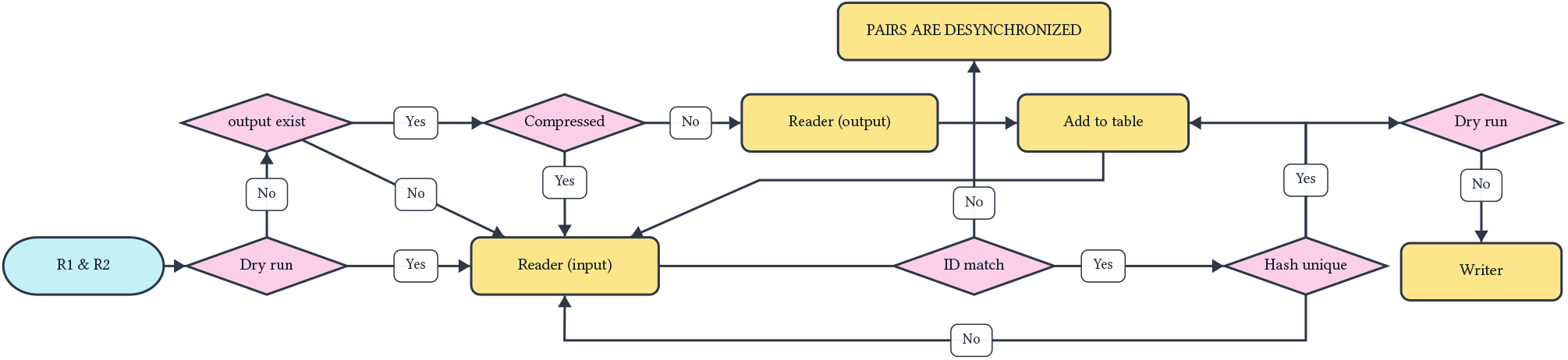
Architectural design of `FastDedup` for Paired-End reads.

`FastDedup` provides users with precise control over the hash collision rate while maintaining high performance. It automatically scales its hashing strategy, seamlessly switching between 64-bit and 128-bit hashes based on the estimated input sequence count and a user-defined collision probability threshold.

Specifically designed for seamless integration into OMICS pipelines, the tool features robust incremental deduplication and auto-recovery mechanisms. In the event of an interruption, `FastDedup` can safely preload existing hashes from the output file to prevent duplicate processing and smoothly resume operations. If an uncompressed output file becomes corrupted due to a crash, the tool automatically detects the issue, calculates a fail-safe truncation point, and truncates the file to the last valid sequence before continuing.

Finally, a dry-run mode (`--dry-run` or `-s`) is available to calculate the duplication rate without writing any output files. Since file I/O operations account for the majority of execution time, this mode is exceptionally fast and serves as a highly efficient feature for pipeline planning and data analyses.

### 3.1: Algorithmic Complexity and Hardware-Level Optimizations

The architectural principle of `FastDedup`’s performance lies in shifting the deduplication problem from a string-matching paradigm to an integer-matching paradigm. Transforming raw paired-end sequences (e.g., 2 × 150 bytes) into a single 64-bit hash reduces the data by 37.5 times in both space and time complexity.

#### 3.1.1 Space Complexity and Memory Footprint

In a deterministic approach, storing a read pair requires retaining sequences of length *L* (where *L* = *r*_1_ + *r*_2_). The space complexity for *N* unique sequences scales linearly as *O*(*N* × *L*). For a standard 150 bp pairedend dataset, each pair consumes at least 300 bytes of raw character data, plus substantial data structure overhead (e.g., string pointers). Storing 10^8^ sequences typically exceeds 32 GB of RAM. By computing a 64-bit hash using `xxh3`, `FastDedup` reduces the sequence identity into exactly 8 bytes. This decouples the memory requirement from the read length, reducing the space complexity to *O*(*N*) (since 8 ≪ *L*).

#### 3.1.2 Time Complexity and Arithmetic Logic Unit (ALU) Efficiency

String-based deduplication necessitates comparing sequences letter by letter, meaning the time it takes scales with the read length. In the worst-case scenario, comparing two sequences requires *O*(*L*) time. `FastDedup` changes this dynamic. While it still takes *O*(*L*) time to read a sequence and calculate its initial hash, every subsequent comparison uses only that hash. Because the hash is a simple 64-bit integer, the processor’s Arithmetic Logic Unit can compare two of them natively in a single clock cycle, instantly dropping the lookup and comparison time complexity to *O*(1).

#### 3.1.3 Cache Locality and Zero-Allocation Pipeline

Beyond theoretical Big-*O* improvements, the 8-byte hash architecture heavily exploits modern CPU hardware design. Processors fetch data from RAM in discrete 64-byte cache lines. A 300-byte sequence spans multiple cache lines, leading to frequent cache misses and severe memory bandwidth bottlenecks. In contrast, exactly eight 64-bit hashes fit perfectly into a single cache line. As `fxhash` probes the internal hash set for collisions, it operates almost exclusively within the ultra-fast CPU cache (L1/L2), minimizing latency and maximizing throughput.

Furthermore, `FastDedup` processes byte slices directly from the `needletail` [18] parser into the `xxh3` hasher without ever allocating the raw string to the heap. This zero-allocation pipeline completely bypasses system-level memory allocation (`malloc`) overheads, ensuring that the execution time is strictly bound by disk I/O rather than RAM latency.

### 3.2: The Mathematics of Dynamic Hashing

While `FastDedup` defaults to the highly efficient 64-bit hashing strategy of `xxh3`, it dynamically assesses the risk of hash collisions to guarantee data integrity. The theoretical risk of a hash collision probability *p* for *x* sequences follows the standard Birthday Problem approximation [19,20]:

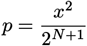

Where *N* ∈ {64, 128} represents the bit-length of the hash. This probability expresses the chance that **at least one collision exists anywhere in the entire dataset**, not a per-read false-duplicate rate. When a collision does occur, it affects exactly one read; since *p* is small, the expected number of false duplicates is also approximately *p*.

At the default 64-bit configuration and a probability threshold of 0.01, it requires approximately 0.6 × 10^9^ sequences before the probability of any single collision occurring reaches 1%. In practical terms, even at this boundary, a run would produce fewer than one spurious duplicate on average. If the estimated dataset size pushes the collision probability beyond this threshold, `FastDedup` automatically scales to 128-bit hashing, extending the same guarantee to approximately 2.6 × 10^18^ sequences — far beyond any current sequencing experiment.

In everyday use cases, the 64-bit hashing strategy is more than sufficient. The 128-bit option serves as a safeguard for extreme cases, ensuring that `FastDedup` can maintain its performance and integrity even as sequencing datasets continue to grow exponentially. Even at the theoretical boundary, the expected number of false duplicates remains below one, and any dataset large enough to push beyond that boundary would contain hundreds of millions of reads, making a handful of removals statistically insignificant relative to the overall data volume.

For paired-end reads, maintaining this theoretical probability requires careful bitwise operations. Let *S*_1_ and *S*_2_ represent the nucleotide sequences of *R*1 and *R*2, and *H* be our non-cryptographic uniform hash function mapping to {0, 1}^*N*^ . We define the independent hashes for each read as:

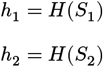

If the algorithm combined these hashes using a simple bitwise XOR (⊕) to create a unified fingerprint *h*_*u*_, a critical failure occurs when *R*1 and *R*2 are identical sequences (*S*_1_ = *S*_2_). In this case, *h*_1_ = *h*_2_, resulting in:

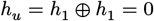

Every paired-end read with identical forward and reverse sequences would map to 0. This would cause massive, deterministic hash collisions, meaning the collision probability would no longer be bounded by the theoretical uniform distribution, but rather by the biological frequency of identical sequence pairs in the dataset.

To prevent this symmetrical cancellation, the algorithm introduces a bitwise left-rotation by *N*/2 bits, denoted as *R*(*x, k*) where *x* is the hash and *k* is the number of bits to rotate.

The unified fingerprint is instead calculated as:

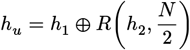

Under this operation, even if *S*_1_ = *S*_2_:

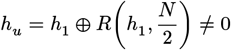

Because 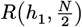 shifts the lower half of the bits to the upper half and vice versa, 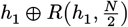 will almost never equal 0. It would only equal 0 if *h*_1_ consisted of two perfectly identical halves, an event with an astronomically low probability of 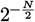.

Since a bitwise rotation is a bijective function over the *N*-bit vector space, it perfectly preserves the original entropy of *h*_2_. When two independent, uniformly distributed random variables are combined via XOR, the resulting variable remains uniformly distributed. Therefore, the unified fingerprint *h*_*u*_ safely maintains a strict uniform distribution, ensuring the overall collision risk remains uncorrupted and strictly bounded by the initial probability formula.

This is the mathematical foundation that allows `FastDedup` to guarantee a collision probability of less than 1% for datasets containing up to approximately 600 million paired-end reads (1.2 billion total sequences) when using 64-bit hashes.

### 3.3: FxHash: Algorithmic Mechanics and ALU Optimizations

While `FastDedup` utilizes `xxh3` to compute sequence fingerprints, it relies on `fxhash` [17] to manage these fingerprints within its internal hash set. Concretely, `FastDedup` uses a `FxHashSet`, a hash set data structure that uses `fxhash` as its internal bucket-routing function: the fixed-size fingerprint produced by `xxh3` is rehashed by `fxhash` to determine its slot in the table. This two-level design is intentional, `xxh3` provides highquality, collision-resistant fingerprints of biological sequences, while `fxhash` provides minimal-overhead addressing into the hash table itself.

`fxhash` is a high-performance, non-cryptographic hashing algorithm derived from the internal hasher utilized in the Rust compiler (`rustc`) and Firefox. Its architectural strength lies not in collision resistance, but rather in raw computational throughput and extremely low latency.

For every word extracted from the input, `fxhash` applies a highly optimized state transition function. Let *H* represent the internal state of the hash (initialized to 0), and *W* represent an incoming data word:

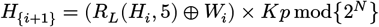

Where:

- *N* is the machine word size (typically 64 bits).
- *R*_*L*_(*x, k*) denotes a bitwise left-rotation of the state *x* by *k* bits (`ROTATE = 5`).
- *K*_*p*_ is a predefined *p* = *N*-bit magic constant (e.g. *K*_64_ = 0x517cc1b727220a95).

The bitwise left-rotation ensures that previously hashed bits propagate rapidly across the state vector. Being a bijective function, it preserves the entropy of *H*_*i*_ while preventing localized clustering. The XOR then introduces the new word *W*_*i*_ in a single clock cycle, and the wrapping multiplication by the large prime-like constant *K* acts as an avalanche multiplier, diffusing the newly mixed entropy across all 64 bits. The algorithm explicitly relies on modulo 2^*N*^ arithmetic, leveraging the CPU’s native handling of integer overflow without any computational penalty.

Within `FastDedup` specifically, `fxhash` receives only the fixed-size fingerprints already produced by `xxh3`, either 64-bit or 128-bit values, corresponding to *L* ∈ {8, 16} bytes. Because *L* is bounded to at most 2 machine words, the state transition function executes in exactly 1 or 2 iterations regardless of the input dataset size, making the bucket-routing step a strict *O*(1) operation. This stands in contrast to the general case, where a byte-oriented hasher processing an arbitrary sequence of length *L* would require *O*(*L*) state transitions.

The algorithm operates entirely in-place on a single internal state variable and requires no auxiliary data structures, lookup tables, or intermediate buffers, yielding an auxiliary space complexity of *O*(1). Combined with the zero-allocation and cache-locality properties described in Section 3.1, this ensures the memory footprint remains flat during hash table operations, preventing fragmentation and avoiding OS-level memory management overhead.

### 3.4: Architecture

## 4 Benchmarking and performance evaluation

To test the performance of the different tools mentioned in the previous sections (Table 3), we benchmarked them on a series of synthetic datasets of varying sizes, ranging from 100,000 to 300 million paired-end reads, to which we added 10% duplicates. These reads were generated based on a real human whole genome (GRCh38). This allows us to evaluate the scalability of each tool and its suitability for different types of sequencing projects, from small-scale experiments to large population genomics studies. To run these benchmarks in a controlled environment, we set up a Snakemake pipeline where each tool had 1 core and 33 GB of RAM available. This pipeline ran on the High-Performance Computing (HPC) ISDM-MESO cluster with AMD EPYC 9654 processors. If a tool exceeds 32 GB of RAM, it is killed and documented as OOM (Out Of Memory) in the results table. At a time when RAM and electricity are becoming increasingly costly and environmentally impactful, it is crucial to consider the resource efficiency of bioinformatics tools. Since these tools represent the first step in data preprocessing, it is inefficient to run them using excessive resources.

**Table 1:** Paired-end duplication rate analysis.

| Tool<br>Duplication (%) | 0.1M | 0.3M | 0.5M | 1M | 3M | 5M | 10M | 30M | 50M | 100M | 300M |
| --- | --- | --- | --- | --- | --- | --- | --- | --- | --- | --- | --- |
| fdedup_h128 | 10.00 | 10.00 | 10.00 | 10.00 | 10.00 | 10.00 | 10.00 | 10.00 | 10.00 | 10.00 | 10.00 |
| fdedup_h64 | 10.00 | 10.00 | 10.00 | 10.00 | 10.00 | 10.00 | 10.00 | 10.00 | 10.00 | 10.00 | 10.00 |
| fastp | 10.00 | 10.00 | 10.00 | 10.00 | 10.00 | 10.00 | 10.00 | 10.00 | 10.00 | 10.00 | 10.00 |
| clumpify | 10.00 | 10.00 | 10.00 | 10.00 | 10.00 | 10.00 | 10.00 | 10.00 | 10.00 | 10.00 | — |
| cdhitdup | 10.00 | 10.00 | 10.00 | 10.00 | 10.00 | 10.00 | 10.00 | — | — | — | — |
| prinseqpp | 10.00 | 10.00 | 10.00 | 10.00 | 10.00 | 10.00 | 10.01 | 23.78 | 48.66 | 74.04 | 91.35 |

**Table 2:**
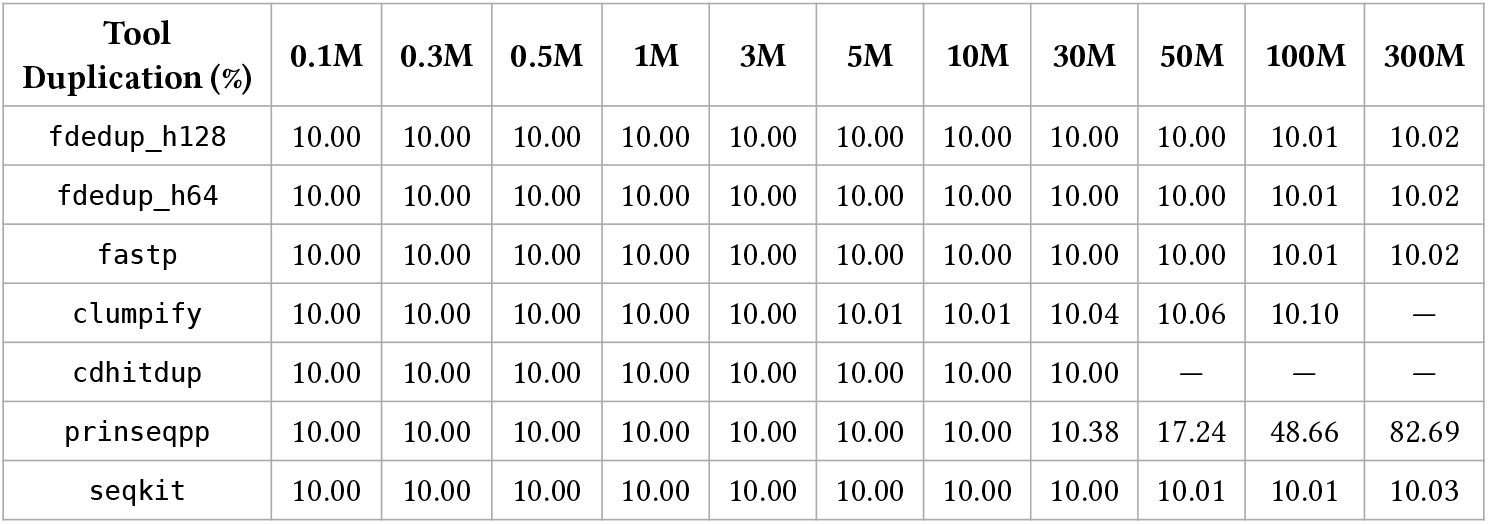
Single-end duplication rate analysis.

| Tool<br>Duplication (%) | 0.1M | 0.3M | 0.5M | 1M | 3M | 5M | 10M | 30M | 50M | 100M | 300M |
| --- | --- | --- | --- | --- | --- | --- | --- | --- | --- | --- | --- |
| fdedup_h128 | 10.00 | 10.00 | 10.00 | 10.00 | 10.00 | 10.00 | 10.00 | 10.00 | 10.00 | 10.01 | 10.02 |
| fdedup_h64 | 10.00 | 10.00 | 10.00 | 10.00 | 10.00 | 10.00 | 10.00 | 10.00 | 10.00 | 10.01 | 10.02 |
| fastp | 10.00 | 10.00 | 10.00 | 10.00 | 10.00 | 10.00 | 10.00 | 10.00 | 10.00 | 10.01 | 10.02 |
| clumpify | 10.00 | 10.00 | 10.00 | 10.00 | 10.00 | 10.01 | 10.01 | 10.04 | 10.06 | 10.10 | — |
| cdhitdup | 10.00 | 10.00 | 10.00 | 10.00 | 10.00 | 10.00 | 10.00 | 10.00 | — | — | — |
| prinseqpp | 10.00 | 10.00 | 10.00 | 10.00 | 10.00 | 10.00 | 10.00 | 10.38 | 17.24 | 48.66 | 82.69 |
| seqkit | 10.00 | 10.00 | 10.00 | 10.00 | 10.00 | 10.00 | 10.00 | 10.00 | 10.01 | 10.01 | 10.03 |

**Table 3:** Summary of existing tools for reads deduplication and their limitations.

| Tool | Algorithm / Approach | PE Support | Error Tolerance | Memory Usage | Key Limitations |
| --- | --- | --- | --- | --- | --- |
| fastp [3,4] | Probabilistic (Bloom filter/hashes) | Yes | No | Configurable | Long runtime. |
| FastUniq [5] | Deterministic (Merge sort) | Yes | No | Very High | Severe RAM bottleneck; sensitive to input file order. |
| clumpify [7] | k-mer grouping | Yes | Yes | Variable | Variable stability on massive datasets; high memory overhead. |
| seqkit rmdup [8,9] | Exact matching (Hashing/MD5) | No | No | Very Low | Desynchronizes paired-end files. |
| CD-HIT-DUP [10,11] | Greedy incremental clustering | Yes | Yes | Very High | Stability issues in containerized environments. |
| prinseq++ [12] | Probabilistic (Bloom filter) | Yes | No | Low | False-positives rate |
| FastDedup | Exact matching (Hashing/xxh3) | Yes | No | Very Low | Theoretical negligible collision risk |

**Table 4:**
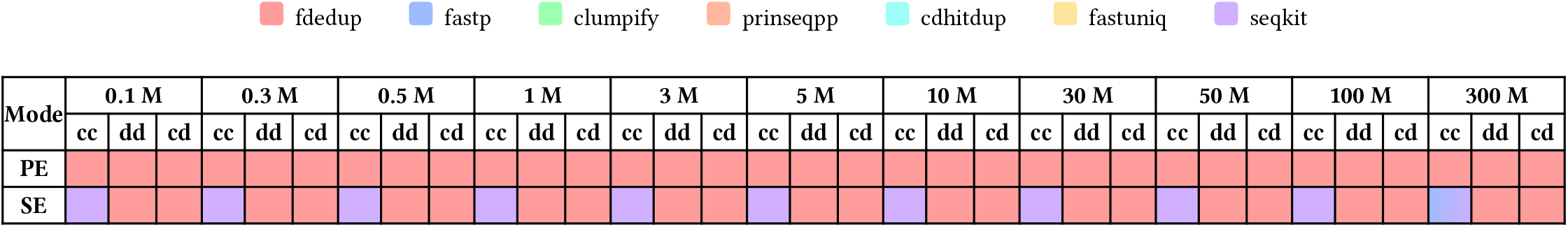
Fastest speed of each tool across different dataset sizes and file formats.

**Table 5:**
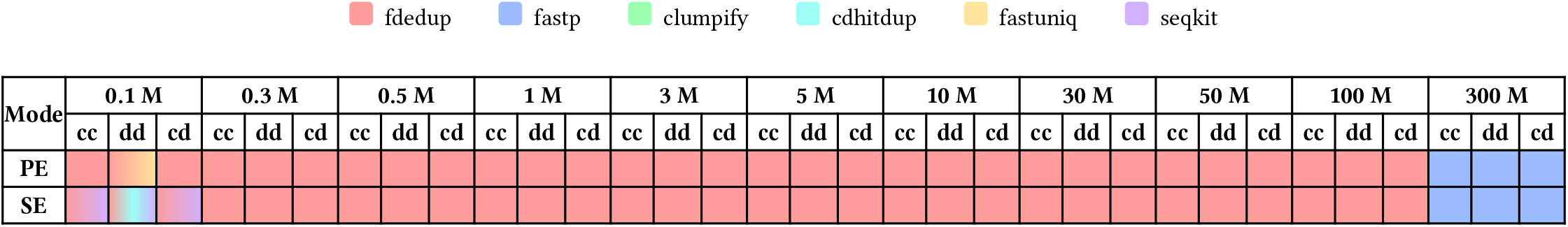
Lowest RAM of each tool across different dataset sizes and file formats. Prinseq++ is excluded from this table because of the duplication rate issue.

### 4.1: Compressed Paired-End FASTQ

Here we will evaluate the performances of the tools in a Paired-End Compressed in Compressed out (PE-cc) scenario. Before discussing the results, it should be noted that `FastUniq` and `CD-HIT-DUP` are excluded from this benchmark because they do not support compressed files. The default compression level of `fastp` is 4, whereas all other tools use a compression level of 6, which is the default for `gzip`.

After setting `fastp` to use the same compression level, we observe that `FastDedup` is around 50% faster than `fastp` while using significantly less RAM up to 300M reads, at which point it uses roughly the same amount as `fastp` for the 64-bit version (Figure 4, Table 6). We also observed no significant differences in execution time between the 64-bit and 128-bit hashing strategies. Furthermore, the RAM usage for the 128-bit strategy is strictly doubled compared to the 64-bit strategy, perfectly reflecting the twofold increase in fingerprint size. Looking closely at the memory scaling, `clumpify`’s RAM usage increases steeply, exhausting the 32 GB limit before reaching 100M reads (consuming over 23 GB at 50M reads alone). In contrast, `FastDedup` maintains a very flat memory profile, using only around 4.6GB of RAM at 300M reads. Execution time for `FastDedup` and other tools scales almost linearly with volume, demonstrating their respective I/O-bound bottlenecks, but `FastDedup` remains faster than `fastp` by approximately 50%.

**Figure 4:**
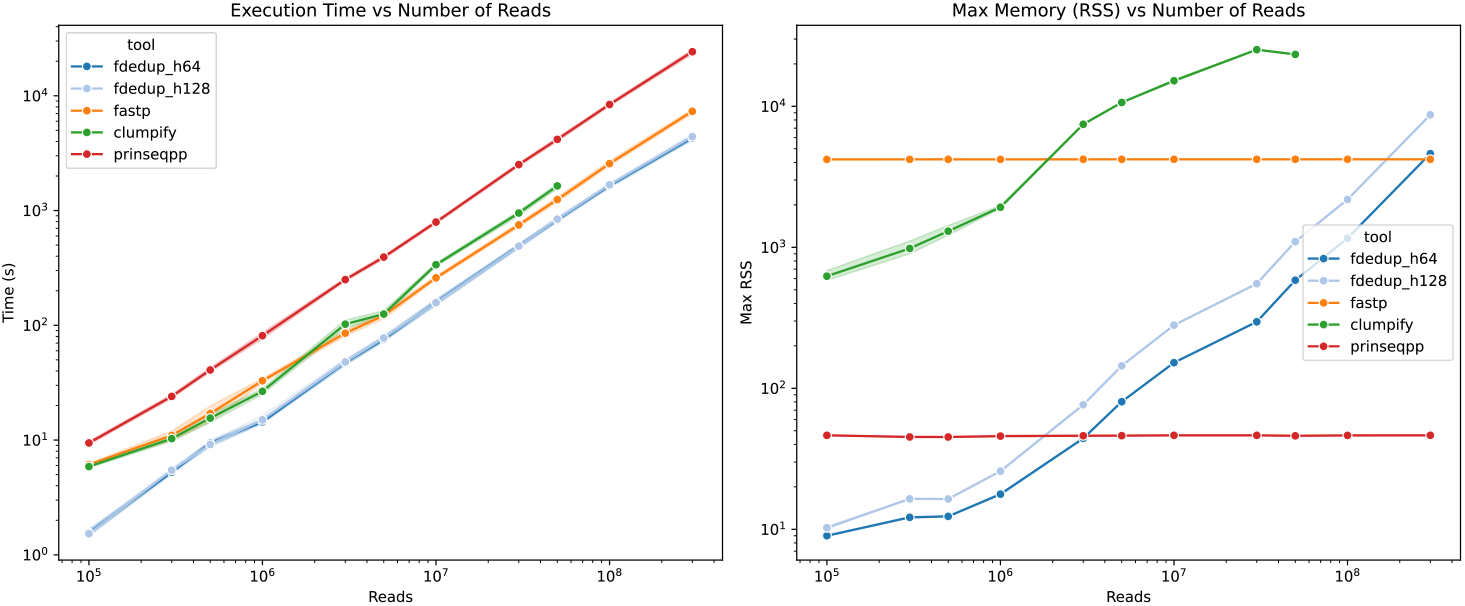
Execution time and memory usage for Compressed Paired-End FASTQ deduplication.

**Table 6:** Execution time and memory usage for compressed input and compressed output paired-end FASTQ files.

| Tool<br>(s)<br>(MB) | 0.1M | 0.3M | 0.5M | 1.0M | 3.0M | 5.0M | 10.0M | 30.0M | 50.0M | 100.0M | 300.0M |
| --- | --- | --- | --- | --- | --- | --- | --- | --- | --- | --- | --- |
| clumpify | 5.9±0.2<br>624±90 | 10.3±1.0<br>982±203 | 15.5±2.1<br>1302±187 | 26.6±2.5<br>1924±75 | 102±14.6<br>7458±137 | 125±14.1<br>10650±15 | 337±21.1<br>15185±494 | 952±78.4<br>25226±42 | 1639±131<br>23350±271 | OOM | OOM |
| fastp | 6.1±0.2<br>4202±0 | 11.0±1.7<br>4205±1 | 17.0±4.5<br>4207±1 | 32.9±2.7<br>4203±1 | 85.5±8.1<br>4208±1 | 122±14.3<br>4208±1 | 259±16.1<br>4209±1 | 750±51.8<br>4210±1 | 1246±108<br>4209±1 | 2564±205<br>4209±1 | 7324±566<br>4210±2 |
| fdedup_h128 | 1.5±0.1<br>10±0 | 5.5±0.4<br>16±0 | 9.2±0.9<br>16±0 | 15.0±2.2<br>26±0 | 47.9±4.5<br>76±0 | 77.5±5.0<br>144±0 | 158±12.8<br>281±0 | 489±33.6<br>552±1 | 841±79.0<br>1096±0 | 1679±95.4<br>2184±0 | 4407±361<br>8712±1 |
| fdedup_h64 | 1.6±0.2<br>9±0 | 5.3±0.4<br>12±0 | 9.4±0.8<br>12±0 | 14.4±1.2<br>18±0 | 47.2±4.6<br>44±0 | 75.4±7.0<br>80±0 | 159±15.9<br>152±0 | 493±36.7<br>296±1 | 830±46.4<br>584±1 | 1649±95.8<br>1160±0 | 4292±340<br>4616±1 |
| prinseqpp | 9.4±0.6<br>46±0 | 24.0±1.5<br>45±0 | 40.8±3.2<br>45±0 | 81.2±8.2<br>46±1 | 250±13.9<br>46±0 | 393±25.2<br>46±0 | 793±41.4<br>46±0 | 2515±154<br>46±0 | 4169±271<br>46±1 | 8399±465<br>46±0 | 24202±1747<br>46±0 |

### 4.2: Uncompressed Paired-End FASTQ

Here we will evaluate the performances of the tools in a Paired-End Uncompressed in Uncompressed out (PE-dd) scenario. For uncompressed FASTQ files as inputs and outputs, the raw computational speed of the tools is fully exposed. `FastDedup` consistently outpaces all other tools in execution time (Figure 5, Table 7), completing 300M reads in under 10 minutes (approx. 450s). In comparison, `prinseq++` requires approximately 2700s and `fastp` roughly 4500s for the same volume—meaning `FastDedup` is around 6× faster than `prinseq+ +` and 10× faster than `fastp`. We can also see that `FastDedup` maintains a highly stable memory footprint (around 4.6GB and 8.7GB of RAM for 64-bit and 128-bit hashing respectively at 300M reads), while `prinseq+ +` edges it out in strict memory efficiency starting at 5M reads. Deterministic string-matching tools struggle significantly with memory capacity here: `CD-HIT-DUP` and `FastUniq` load full sequences into RAM, causing vertical spikes in their memory usage leading to OOM crashes early on (at 30M and 50M reads respectively). `clumpify` also succumbed to OOM errors at 300M reads.

**Figure 5:**
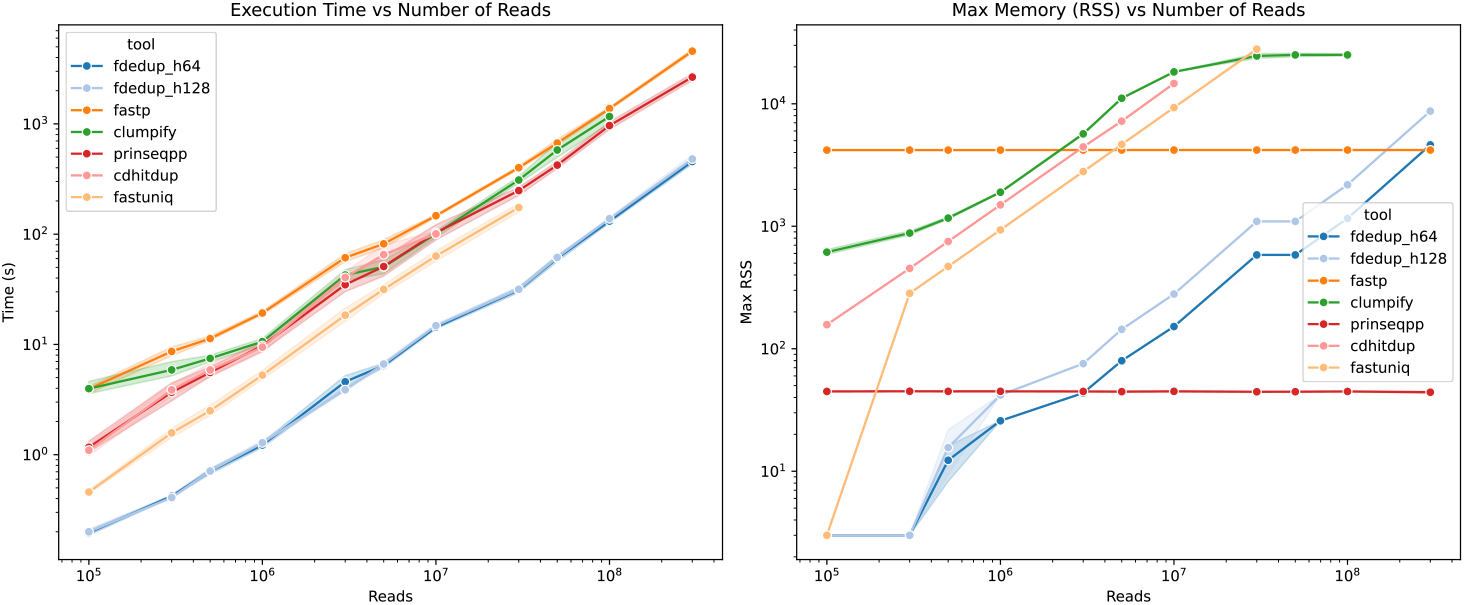
Execution time and memory usage for Uncompressed Paired-End FASTQ deduplication.

**Table 7:** Execution time and memory usage for uncompressed input and uncompressed output paired-end FASTQ files.

| **Tool**<br>(s)<br>(MB) | **0.1M** | **0.3M** | **0.5M** | **1.0M** | **3.0M** | **5.0M** | **10.0M** | **30.0M** | **50.0M** | **100.0M** | **300.0M** |
| --- | --- | --- | --- | --- | --- | --- | --- | --- | --- | --- | --- |
| cdhitdup | 1.1±0.2<br>157±1 | 3.9±0.8<br>453±0 | 5.9±1.4<br>751±0 | 9.4±1.6<br>1495±1 | 40.4±8.9<br>4443±1 | 65.2±23.7<br>7214±125 | 101±11.0<br>14657±39 | OOM | OOM | OOM | OOM |
| clumpify | 4.0±0.9<br>615±50 | 5.9±1.6<br>880±63 | 7.5±0.9<br>1166±47 | 10.6±1.3<br>1896±36 | 42.6±8.2<br>5683±151 | 50.6±13.5<br>11092±158 | 101±12.3<br>18211±37 | 310±35.3<br>24576±1982 | 580±155<br>25061±1379 | 1167±303<br>25057±791 | OOM |
| fastp | 3.9±0.5<br>4187±0 | 8.6±1.4<br>4187±1 | 11.3±1.0<br>4188±1 | 19.3±1.2<br>4188±1 | 61.2±9.0<br>4191±0 | 81.7±11.7<br>4192±0 | 147±7.9<br>4192±1 | 401±21.7<br>4192±0 | 670±82.4<br>4192±0 | 1378±75.7<br>4192±2 | 4569±336<br>4192±1 |
| fastuniq | 0.5±0.0<br>3±0 | 1.6±0.3<br>284±0 | 2.5±0.5<br>470±0 | 5.3±0.9<br>935±0 | 18.4±4.8<br>2796±0 | 31.6±5.0<br>4658±0 | 63.3±11.5<br>9312±0 | 174±27.6<br>27927±0 | OOM | OOM | OOM |
| fdedup_h128 | 0.2±0.0<br>3±0 | 0.4±0.0<br>3±0 | 0.7±0.1<br>16±11 | 1.3±0.1<br>42±0 | 3.9±0.2<br>76±0 | 6.6±0.6<br>144±0 | 14.8±1.0<br>280±1 | 31.6±2.9<br>1096±0 | 61.5±4.9<br>1095±0 | 138±6.4<br>2184±0 | 481±76.6<br>8712±0 |
| fdedup_h64 | 0.2±0.0<br>3±0 | 0.4±0.0<br>3±0 | 0.7±0.1<br>12±6 | 1.2±0.1<br>26±0 | 4.6±0.9<br>44±0 | 6.4±0.5<br>80±0 | 14.3±0.9<br>152±0 | 31.1±3.2<br>584±1 | 60.4±6.9<br>584±0 | 131±9.1<br>1159±0 | 456±50.0<br>4616±0 |
| prinseqpp | 1.2±0.2<br>45±0 | 3.7±1.2<br>45±0 | 5.6±1.0<br>45±0 | 9.7±1.6<br>45±0 | 34.8±9.1<br>45±0 | 50.9±16.7<br>45±0 | 103±29.2<br>45±0 | 248±36.7<br>45±0 | 423±56.2<br>45±0 | 965±128<br>45±0 | 2658±326<br>44±1 |

### 4.3: Compressed in, uncompressed out Paired-End FASTQ

Here we will evaluate the performances of the tools in a Paired-End Compressed in Uncompressed out (PE-cd) scenario.

We observe very similar results to those obtained with uncompressed files (Figure 6, Table 8). `FastDedup` being the fastest tool and using the least amount of RAM up until 5M reads, where `prinseq++` starts to use less memory than `FastDedup`. We can also note very similar results for relative performance between each tool, which means that all the tools scale in a similar way whether the input files are compressed or not.

**Figure 6:**
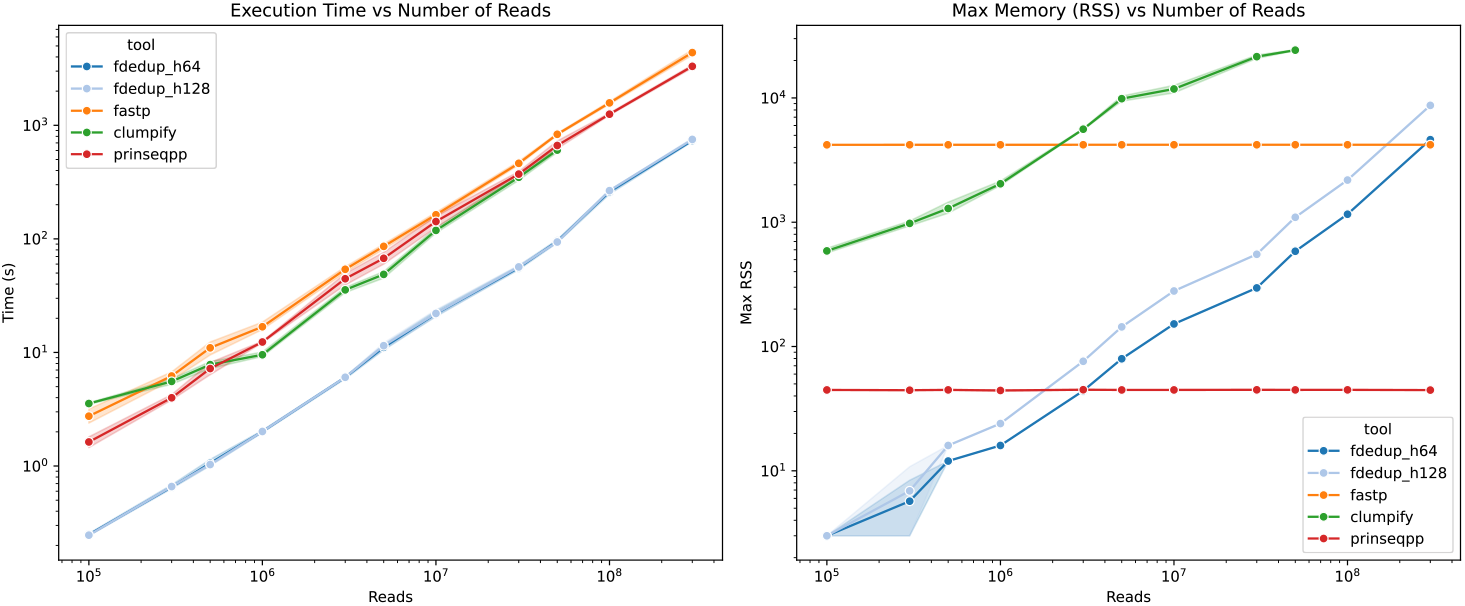
Execution time and memory usage for Compressed-input to Uncompressed-output Paired-End FASTQ deduplication.

**Table 8:** Execution time and memory usage for compressed input and uncompressed output paired-end FASTQ files.

| **Tool**<br>(s)<br>(MB) | **0.1M** | **0.3M** | **0.5M** | **1.0M** | **3.0M** | **5.0M** | **10.0M** | **30.0M** | **50.0M** | **100.0M** | **300.0M** |
| --- | --- | --- | --- | --- | --- | --- | --- | --- | --- | --- | --- |
| clumpify | 3.6±0.1<br>588±45 | 5.6±0.5<br>979±77 | 7.8±0.9<br>1290±224 | 9.5±0.8<br>2039±168 | 35.6±3.2<br>5604±99 | 48.7±5.3<br>9863±796 | 119±10.5<br>11807±1392 | 349±41.8<br>21482±1170 | 605±50.8<br>24234±364 | OOM | OOM |
| fastp | 2.7±0.7<br>4196±1 | 6.2±0.9<br>4197±1 | 11.0±2.4<br>4197±1 | 16.8±2.6<br>4197±1 | 54.0±5.0<br>4200±0 | 85.9±7.3<br>4200±0 | 163±13.6<br>4200±1 | 463±30.4<br>4200±0 | 834±32.3<br>4200±1 | 1576±74.8<br>4200±1 | 4369±421<br>4201±1 |
| fdedup_h128 | 0.2±0.0<br>3±0 | 0.7±0.0<br>7±6 | 1.0±0.0<br>16±0 | 2.0±0.1<br>24±0 | 6.0±0.1<br>76±0 | 11.5±1.2<br>144±0 | 22.1±2.7<br>280±1 | 56.6±3.3<br>552±0 | 93.9±3.8<br>1096±1 | 266±5.0<br>2184±0 | 751±34.7<br>8712±0 |
| fdedup_h64 | 0.2±0.0<br>3±0 | 0.7±0.0<br>6±4 | 1.1±0.1<br>12±0 | 2.0±0.1<br>16±0 | 6.0±0.3<br>44±0 | 11.1±1.1<br>80±0 | 21.9±2.1<br>152±0 | 55.7±3.8<br>296±0 | 95.3±5.2<br>584±0 | 260±17.9<br>1160±0 | 734±29.6<br>4615±0 |
| prinseqpp | 1.6±0.3<br>45±0 | 4.0±0.4<br>45±0 | 7.2±1.6<br>45±0 | 12.4±0.3<br>44±1 | 44.6±8.8<br>45±0 | 67.5±14.0<br>45±0 | 142±28.7<br>45±0 | 372±33.7<br>45±0 | 664±101<br>45±0 | 1253±46.9<br>45±0 | 3306±170<br>45±0 |

### 4.4: Compressed Single-End FASTQ

Here we will evaluate the performances of the tools in a Single-End Compressed in Compressed out (SE-cc) scenario.

For compressed single-end data, `SeqKit` and `fastp` hold a slight advantage in execution time (Figure 7, Table 9) over `FastDedup` due to highly optimized compression pipelines. However, pulling attention to the memory graph reveals a dramatic difference in scalability: while `SeqKit`’s memory scales aggressively up to approximately 19 GB at 300M reads, `FastDedup`’s memory footprint strictly remains stable around 4.6 GB (a flat fourfold reduction). This showcases `FastDedup`’s superior suitability for memory-restricted environments, even when it is roughly 25% slower in pure elapsed time.

**Figure 7:**
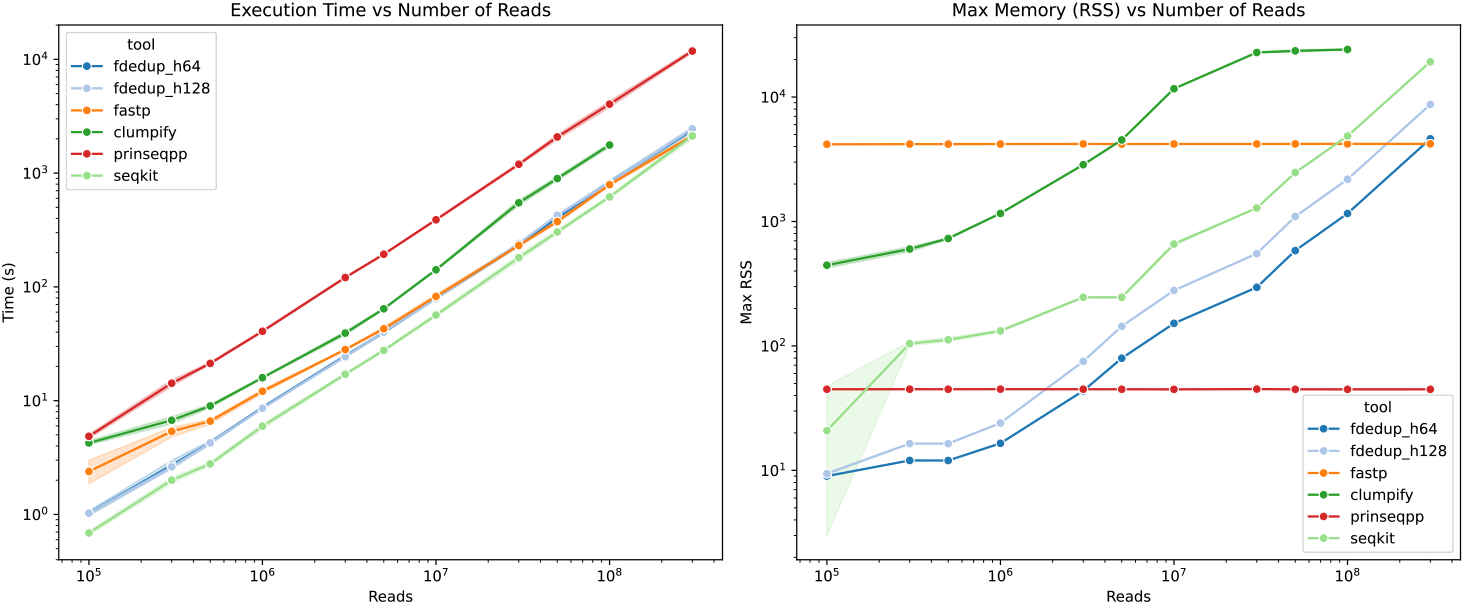
Execution time and memory usage for Compressed Single-End FASTQ deduplication.

**Table 9:**
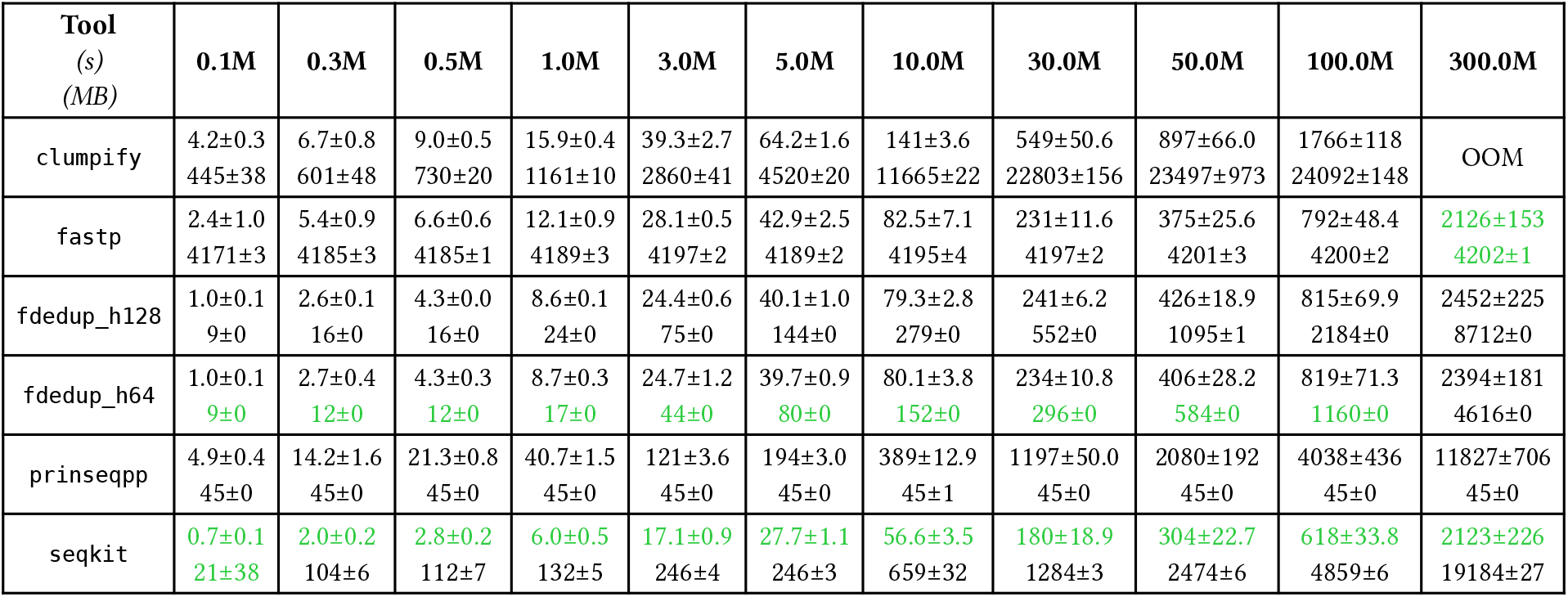
Execution time and memory usage for compressed input and compressed output single-end FASTQ files.

| **Tool**<br>(s)<br>(MB) | **0.1M** | **0.3M** | **0.5M** | **1.0M** | **3.0M** | **5.0M** | **10.0M** | **30.0M** | **50.0M** | **100.0M** | **300.0M** |
| --- | --- | --- | --- | --- | --- | --- | --- | --- | --- | --- | --- |
| clumpify | 4.2±0.3<br>445±38 | 6.7±0.8<br>601±48 | 9.0±0.5<br>730±20 | 15.9±0.4<br>1161±10 | 39.3±2.7<br>2860±41 | 64.2±1.6<br>4520±20 | 141±3.6<br>11665±22 | 549±50.6<br>22803±156 | 897±66.0<br>23497±973 | 1766±118<br>24092±148 | OOM |
| fastp | 2.4±1.0<br>4171±3 | 5.4±0.9<br>4185±3 | 6.6±0.6<br>4185±1 | 12.1±0.9<br>4189±3 | 28.1±0.5<br>4197±2 | 42.9±2.5<br>4189±2 | 82.5±7.1<br>4195±4 | 231±11.6<br>4197±2 | 375±25.6<br>4201±3 | 792±48.4<br>4200±2 | 2126±153<br>4202±1 |
| fdedup_h128 | 1.0±0.1<br>9±0 | 2.6±0.1<br>16±0 | 4.3±0.0<br>16±0 | 8.6±0.1<br>24±0 | 24.4±0.6<br>75±0 | 40.1±1.0<br>144±0 | 79.3±2.8<br>279±0 | 241±6.2<br>552±0 | 426±18.9<br>1095±1 | 815±69.9<br>2184±0 | 2452±225<br>8712±0 |
| fdedup_h64 | 1.0±0.1<br>9±0 | 2.7±0.4<br>12±0 | 4.3±0.3<br>12±0 | 8.7±0.3<br>17±0 | 24.7±1.2<br>44±0 | 39.7±0.9<br>80±0 | 80.1±3.8<br>152±0 | 234±10.8<br>296±0 | 406±28.2<br>584±0 | 819±71.3<br>1160±0 | 2394±181<br>4616±0 |
| prinseqpp | 4.9±0.4<br>45±0 | 14.2±1.6<br>45±0 | 21.3±0.8<br>45±0 | 40.7±1.5<br>45±0 | 121±3.6<br>45±0 | 194±3.0<br>45±0 | 389±12.9<br>45±1 | 1197±50.0<br>45±0 | 2080±192<br>45±0 | 4038±436<br>45±0 | 11827±706<br>45±0 |
| seqkit | 0.7±0.1<br>21±38 | 2.0±0.2<br>104±6 | 2.8±0.2<br>112±7 | 6.0±0.5<br>132±5 | 17.1±0.9<br>246±4 | 27.7±1.1<br>246±3 | 56.6±3.5<br>659±32 | 180±18.9<br>1284±3 | 304±22.7<br>2474±6 | 618±33.8<br>4859±6 | 2123±226<br>19184±27 |

### 4.5: Uncompressed Single-End FASTQ

Here we will evaluate the performances of the tools in a Single-End Uncompressed in Uncompressed out (SE-dd) scenario.

Removing the `gzip` bottleneck entirely, `FastDedup` processes 300M single-end reads in roughly 363 seconds, demonstrating its fundamental algorithmic advantage in parsing and deduplicating (Figure 8, Table 10). It stands around 3 times faster than both `SeqKit` and `fastp` (which hover around 900-1500s). In addition to speed, it uses a quarter of the RAM of `SeqKit` (4.6 GB vs 19 GB) and roughly the same amount as `fastp`.

**Figure 8:**
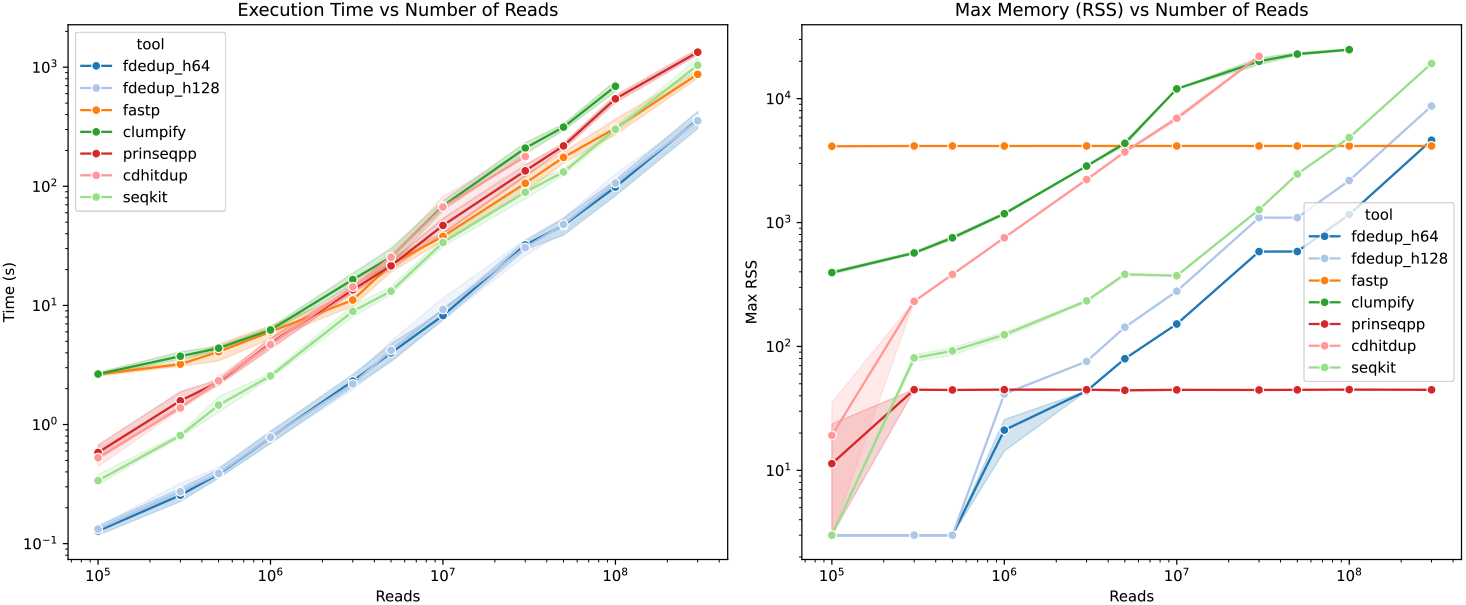
Execution time and memory usage for Uncompressed Single-End FASTQ deduplication.

**Table 10:** Execution time and memory usage for uncompressed input and uncompressed output single-end FASTQ files.

| **Tool**<br>(s)<br>(MB) | **0.1M** | **0.3M** | **0.5M** | **1.0M** | **3.0M** | **5.0M** | **10.0M** | **30.0M** | **50.0M** | **100.0M** | **300.0M** |
| --- | --- | --- | --- | --- | --- | --- | --- | --- | --- | --- | --- |
| cdhitdup | 0.5±0.2<br>19±34 | 1.4±0.1<br>231±0 | 2.3±0.3<br>381±0 | 4.7±0.8<br>754±1 | 14.3±1.9<br>2223±1 | 25.4±3.7<br>3714±1 | 67.0±24.9<br>6943±495 | 178±39.9<br>21958±242 | OOM | OOM | OOM |
| clumpify | 2.7±0.1<br>395±19 | 3.7±0.5<br>568±19 | 4.4±0.3<br>753±39 | 6.2±0.5<br>1179±47 | 16.5±3.3<br>2858±37 | 25.4±7.1<br>4360±309 | 68.5±11.2<br>11985±32 | 209±39.8<br>20013±2244 | 314±33.0<br>22873±853 | 692±122<br>24829±494 | OOM |
| fastp | 2.6±0.1<br>4125±30 | 3.2±0.2<br>4147±1 | 4.1±1.0<br>4147±1 | 6.0±1.2<br>4147±1 | 11.1±2.4<br>4150±1 | 22.1±5.5<br>4150±1 | 38.0±5.0<br>4149±0 | 106±29.1<br>4149±1 | 175±32.0<br>4149±0 | 308±77.5<br>4149±1 | 873±123<br>4150±1 |
| fdedup_h128 | 0.1±0.0<br>3±0 | 0.3±0.1<br>3±0 | 0.4±0.1<br>3±0 | 0.8±0.1<br>41±1 | 2.2±0.4<br>76±0 | 4.2±1.3<br>143±1 | 9.3±3.4<br>279±0 | 30.5±8.4<br>1095±0 | 47.9±11.1<br>1095±0 | 106±34.6<br>2183±1 | 355±49.6<br>8711±1 |
| fdedup_h64 | 0.1±0.0<br>3±0 | 0.3±0.1<br>3±0 | 0.4±0.1<br>3±0 | 0.8±0.2<br>21±10 | 2.3±0.4<br>44±0 | 4.0±1.2<br>80±0 | 8.2±1.3<br>152±0 | 32.2±4.7<br>583±0 | 47.0±12.8<br>583±0 | 98.6±25.1<br>1159±0 | 363±101<br>4615±0 |
| prinseqpp | 0.6±0.1<br>11±18 | 1.6±0.4<br>45±0 | 2.3±0.2<br>45±1 | 4.8±0.9<br>45±0 | 13.5±1.3<br>45±0 | 21.5±2.8<br>44±1 | 47.0±11.4<br>45±0 | 135±27.1<br>45±0 | 218±19.7<br>45±0 | 543±67.5<br>45±0 | 1337±130<br>45±0 |
| seqkit | 0.3±0.1<br>3±0 | 0.8±0.0<br>81±9 | 1.5±0.4<br>92±11 | 2.6±0.2<br>124±10 | 8.9±1.8<br>233±8 | 13.2±1.7<br>382±12 | 33.8±4.8<br>372±6 | 89.3±22.2<br>1271±11 | 132±14.3<br>2463±8 | 301±31.4<br>4848±10 | 1040±224<br>19206±29 |

### 4.6: Compressed in, uncompressed out Single-End FASTQ

Here we will evaluate the performances of the tools in a Single-End Compressed in Uncompressed out (SE-cd) scenario. We got very similar results to those for uncompressed single-end data, with `FastDedup` being the fastest tool for all volumes and using the least amount of RAM up until 5M reads, where `prinseq++` starts to use less memory than `FastDedup` (Figure 9, Table 11).

**Figure 9:**
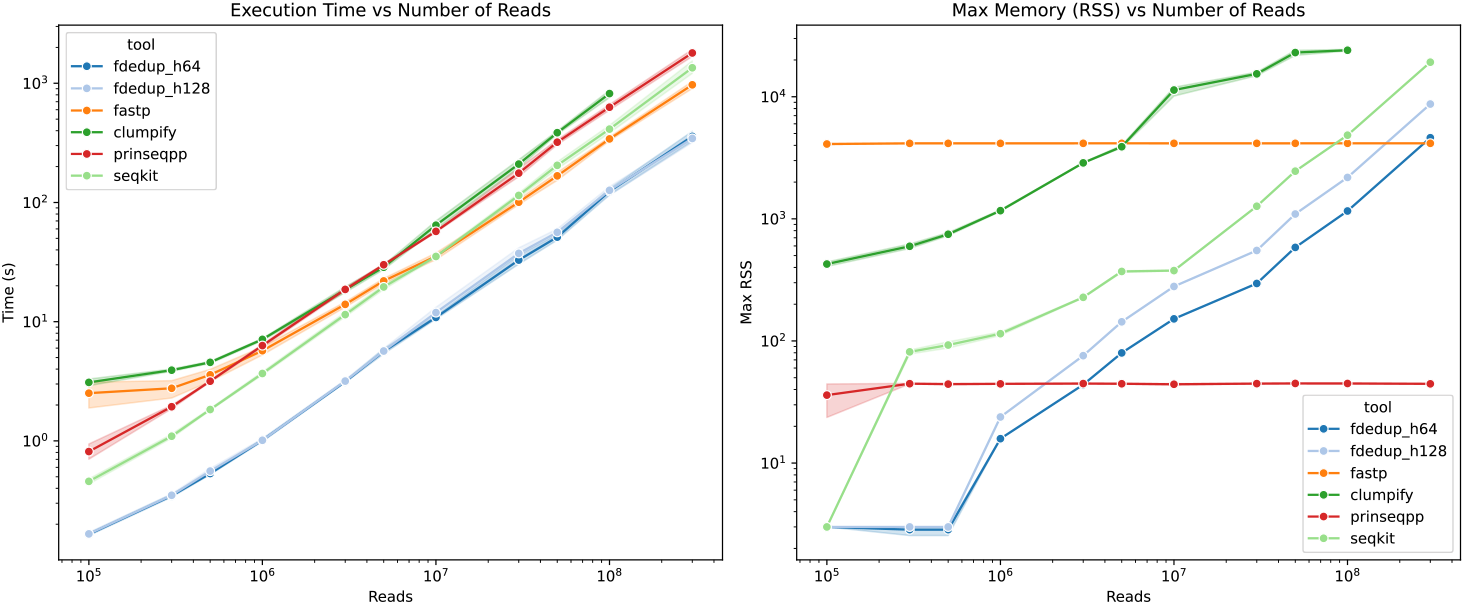
Execution time and memory usage for Compressed-input to Uncompressed-output Single-End FASTQ deduplication.

**Table 11:** Execution time and memory usage for compressed input and uncompressed output single-end FASTQ files.

| **Tool**<br>**(s)**<br>**(MB)** | **0.1M** | **0.3M** | **0.5M** | **1.0M** | **3.0M** | **5.0M** | **10.0M** | **30.0M** | **50.0M** | **100.0M** | **300.0M** |
| --- | --- | --- | --- | --- | --- | --- | --- | --- | --- | --- | --- |
| clumpify | 3.1±0.3<br>426±32 | 3.9±0.1<br>596±45 | 4.6±0.2<br>748±39 | 7.1±0.3<br>1167±29 | 18.5±1.7<br>2873±30 | 28.6±2.3<br>3896±159 | 64.5±9.3<br>11344±1919 | 210±28.0<br>15394±883 | 385±22.3<br>22998±1852 | 820±98.9<br>24023±608 | OOM |
| fastp | 2.5±1.1<br>4100±74 | 2.8±0.8<br>4151±1 | 3.6±0.7<br>4151±0 | 5.7±0.8<br>4151±1 | 14.0±1.2<br>4154±1 | 22.0±1.9<br>4154±1 | 35.4±4.0<br>4153±1 | 101±9.4<br>4153±0 | 167±22.4<br>4153±1 | 341±21.1<br>4154±2 | 971±114<br>4154±2 |
| fdedup_h128 | 0.2±0.0<br>3±0 | 0.3±0.0<br>3±0 | 0.6±0.0<br>3±0 | 1.0±0.0<br>24±0 | 3.2±0.1<br>76±0 | 5.7±0.5<br>143±1 | 11.9±2.0<br>279±0 | 37.4±6.9<br>551±1 | 56.2±6.7<br>1095±1 | 126±17.9<br>2183±1 | 344±45.2<br>8711±1 |
| fdedup_h64 | 0.2±0.0<br>3±0 | 0.3±0.0<br>3±0 | 0.5±0.0<br>3±0 | 1.0±0.0<br>16±0 | 3.1±0.1<br>44±0 | 5.6±0.5<br>80±0 | 10.9±0.7<br>152±0 | 32.9±5.3<br>296±0 | 51.2±4.6<br>584±0 | 124±12.8<br>1159±0 | 358±61.4<br>4616±0 |
| prinseqpp | 0.8±0.2<br>36±17 | 1.9±0.1<br>45±0 | 3.2±0.1<br>44±1 | 6.3±0.3<br>45±0 | 18.7±1.1<br>45±0 | 30.0±1.0<br>45±1 | 57.2±1.6<br>44±1 | 176±26.2<br>45±0 | 321±26.8<br>45±0 | 630±58.4<br>45±0 | 1796±218<br>45±0 |
| seqkit | 0.5±0.0<br>3±0 | 1.1±0.1<br>81±6 | 1.8±0.1<br>93±10 | 3.7±0.2<br>115±7 | 11.5±0.9<br>228±3 | 19.6±1.4<br>371±10 | 35.2±1.5<br>377±5 | 115±8.4<br>1268±4 | 205±23.8<br>2457±6 | 413±53.3<br>4843±7 | 1349±268<br>19182±34 |

## 5 Discussion and future perspectives

Now that we have significant data from isolated benchmarks, we can discuss the implications of these results for real-world applications and the future development of `FastDedup`.

### 5.1: Output Quality and Deduplication Accuracy

Despite each tool employing very different approaches to deduplication, they all produced nearly identical outputs regarding the number of unique sequences and the duplication rate. The differences between tools are marginal, staying entirely within a 0.1% margin. However, we were unable to estimate this metric for `CD-HIT-DUP` because it does not output a log. Moreover, a critical problem emerges with `prinseq++` starting at 10M reads: its reported duplication rate exceeds the expected 10.1%, climbing to over 90% at 300M reads (Figure 10, Table 1). Because of this severe issue, we strongly advise against using `prinseq++` in production pipelines; although it becomes more memory-efficient than `FastDedup` at 5M reads, its false-positive rate becomes unacceptably high by 10M reads.

**Figure 10:**
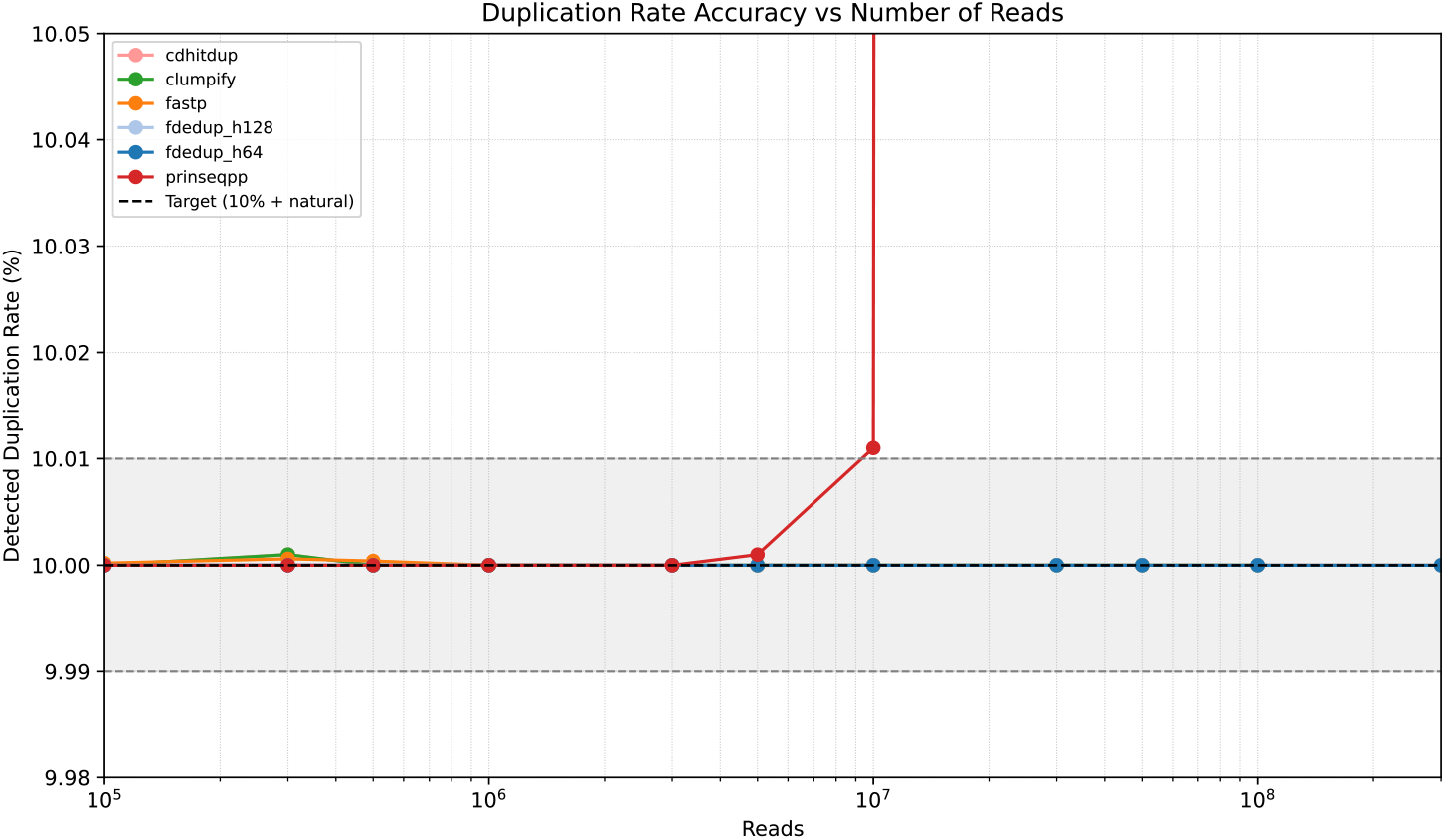
Duplication rate observed for Uncompressed Paired-End FASTQ deduplication.

A similar pattern is observed for `clumpify` on single-end data, but starting at 3M reads. It rises above 10.01% at 10M reads, continuing to climb steadily up to 100M reads. While this still exceeds the false-positive threshold, it remains significantly better than the 90% false-positive rate exhibited by `prinseq++` at 300M reads (Figure 11, Table 2).

**Figure 11:**
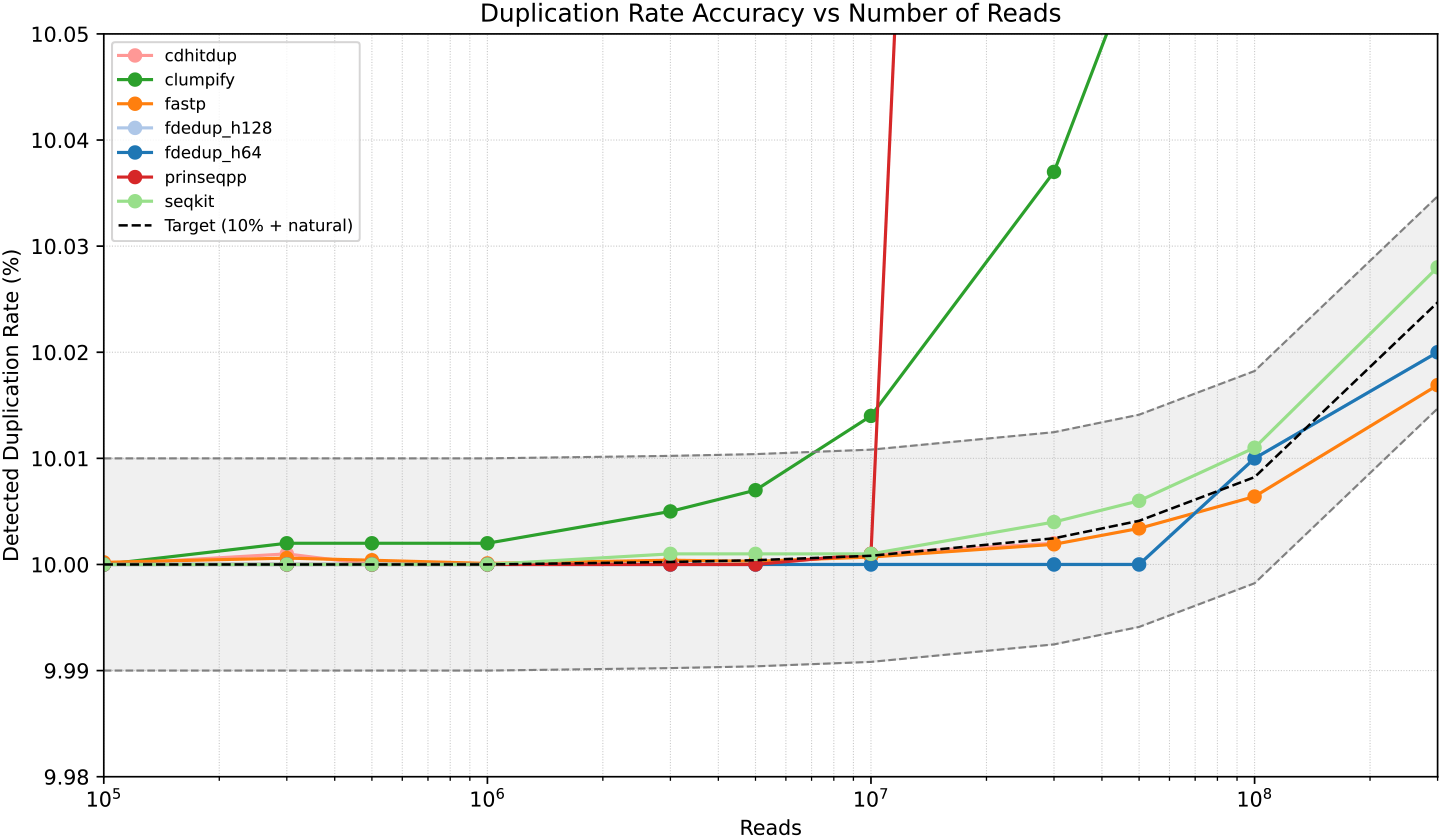
Duplication rate observed for Uncompressed Single-End FASTQ deduplication.

The actual duplication rate is not strictly constant across different sequencing volumes because `wgsim` acts as a random read simulator. We can calculate the expected number of natural coincidental duplicates *C* with *N* randomly generated reads from a uniform distribution of *M* possible start positions as 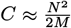. However, the human genome is not uniform, as it contains high-frequency repeat families [21], so we have to approximate *M* with the number of unique start positions in the simulated dataset. We can approximate *k* = (2*G*)^−1^ using a known data point from your `wgsim` simulation. Looking at the generation of 270, 000, 000 single-end reads, the simulation naturally produced roughly 60, 000 coincidental duplicates.

We can solve *k* like:

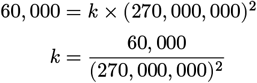

We can now express the number of natural duplicates *C* as *C* = *k* × *N* ^2^ and add this value to the artificial 10% duplication rate we intentionally introduced, yielding the theoretical expected duplication rate for each dataset size.

Before leaving this, we have to talk about why `prinseq++`’s false-positive rate explodes so dramatically with increasing dataset size. We can calculate the expected false-positive rate *F* for a Bloom filter with *m* bits and *n* inserted elements using *k* hash functions as:

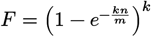

We have multiple data points for `prinseq++`’s false-positive rate at different volumes, so we can solve for *m* using the observed false-positive rates like so:

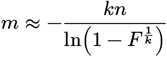

However, we also need *k*, the number of hash functions used by `prinseq++`, which is not documented.

Fortunately, this can be determined by inspecting the code. In `main.cpp`, we can find this snippet of code starting at line 497.

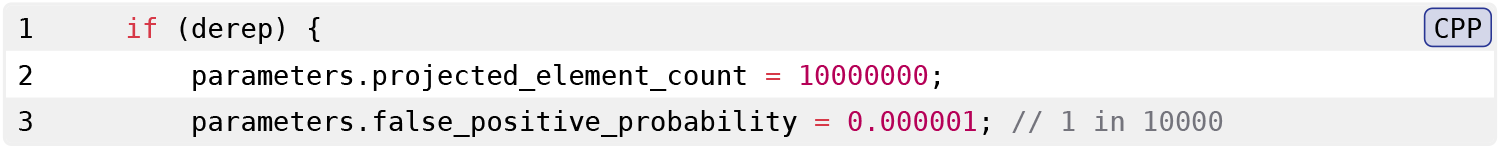

The algorithm begins with two fixed, hardcoded constraints:

- *n* (Projected elements): 10, 000, 000
- *p* (Target false positive probability): 10^−6^

Because a Bloom filter must use a whole number of hash functions, the algorithm performs a discrete optimization. It iterates over integers *k* ∈ ℤ^+^ to find the argument *k* that minimizes the size *m*(*k*):

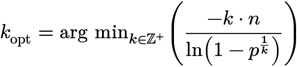

The continuous global minimum for this function occurs when 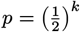, we can directly solve for the ideal *k*:

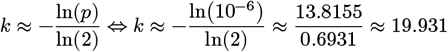

When evaluating the function *m*(*k*) across integer values, the discrete minimum strictly aligns with the closest integer to this theoretical continuous minimum. Therefore, the search loop locks in on: *k*_opt_ = 20 With the optimal number of hash functions established, the algorithm substitutes this value back into the objective function to find the raw bit size:

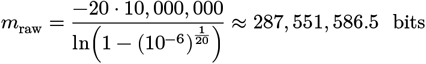

The code then uses `static_cast` to truncate the decimal, resulting in an integer raw size of **287,551**,**586 bits**. We saw experimentally that `prinseq++` had a 45MB RAM footprint at 5M reads, which is consistent with this theoretical calculation since 287, 551, 586 bits is approximately 34.3 MB. The rest of the RAM usage can be attributed to the overhead of the Bloom filter data structure.

Now that we have explicitly determined the internal parameters, we can mathematically model how the false-positive rate degrades when the filter is fed a dataset of 100 million paired-end reads.

Crucially, because `prinseq++` was executed using exclusively the `-derep` flag, no prior quality trimming or length filtering occurred. This guarantees that 100% of the dataset, all 200 million individual sequences, was forced into the Bloom filter. Assuming an estimated true biological duplication rate of 10%, the approximate number of unique sequences inserted into the filter is *N* ≈ 180, 000, 000.

When `prinseq++` outputs the final deduplication percentage, it reports the fraction of reads flagged as duplicates across the entire run. As the filter fills up, the instantaneous false-positive rate after *x* unique elements have been inserted, *F* (*x*), increases monotonically toward saturation:

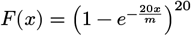

The expected average false-positive rate 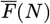 for a dataset containing *N* unique elements is the integral of this instantaneous rate over the domain of *N*, divided by *N*:

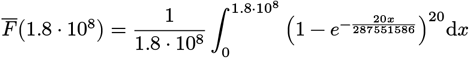

Evaluating this integral yields the theoretical average false-positive rate among unique reads:

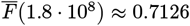

This indicates that 71.26% of the unique reads are expected to be incorrectly flagged as duplicates due to filter saturation. To obtain the total reported duplication rate, we must account for both the true duplicates and the false positives relative to the total number of reads:

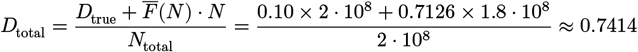

This theoretical prediction of 74.14% is in strikingly close of the empirical observation of 74.04%, differing by only 0.1 percentage points. The exclusive use of the `-derep` flag is what allows this mathematical model to map so cleanly to reality, as there are no “lost” reads from upstream filters skewing the input volume *N*. It is worth noting that this model assumes all *N* unique reads are sequentially inserted into the filter. In practice, reads that trigger a false positive may not advance the filter state, which would slow saturation. The close agreement between the theoretical and empirical values suggests this effect is negligible at this scale. Regardless, the formulation strongly demonstrates that the 34.3 MB hardcoded memory footprint guarantees a catastrophic failure for a 200-million-read insertion volume.

### 5.2: In short

## 6 Conclusion and Recommendations

`FastDedup` represents a significant improvement for FASTX deduplication, combining unmatched speed with a remarkably low memory footprint across all tested scenarios. Its design, shifting the deduplication problem from string-matching to integer-matching through non-cryptographic hashing, allows it to scale from small experimental datasets to the largest population-scale whole-genome sequencing projects without changing configuration or requiring HPC-class hardware.

### 6.1: Summary of findings

For paired-end data, `FastDedup` is the clear winner on both execution time and RAM usage. It consistently outperforms all competitors in execution time, being up to 6× faster than `prinseq++` and more than 10× faster than `fastp` on uncompressed 300M-read datasets, while keeping its memory footprint well below that of every other tool that survived to large volumes except for `fastp` at 300M reads. `FastUniq` and `CD-HIT-DUP` are eliminated by memory constraints before reaching 100M reads, and `clumpify` crashes at 100M reads under the 32GB RAM budget regardless of format for paired-end cc and cd. `prinseq++`, despite being the most memory-efficient competitor, disqualifies itself on correctness grounds: its Bloom filter false-positive rate explodes beyond 10% starting at 10M reads, reaching above 90% at 300M reads, making it unsuitable for any analysis where duplicate removal must be reliable. For single-end data, the picture is more nuanced.

`SeqKit` outperforms `FastDedup` on speed when both input and output are compressed, owing to its mature multi-threaded compression pipeline. However, `FastDedup` reverses this advantage completely on uncompressed I/O, where it runs around 3 times faster than both `SeqKit` and `fastp` at 300M reads, and maintains a drastically lower memory footprint throughout: at 300M reads, `SeqKit` consumes approximately 19 GB of RAM compared to `FastDedup`’s 4.6 GB, a fourfold difference. Users working on memory-constrained nodes or seeking maximum throughput through uncompressed intermediate files should therefore prefer `FastDedup` even for single-end libraries.

### 6.2: Practical recommendations

For maximum performance in any pipeline, we recommend the following strategy: run `FastDedup` immediately after quality control, and direct its output to an uncompressed temporary file. Writing uncompressed output eliminates the compression bottleneck that otherwise dominates execution time, and the temporary files can be recompressed or deleted once downstream steps are complete. For users whose storage constraints require compressed outputs throughout, the compressed-output mode of `FastDedup` remains faster than all alternatives for paired-end data, and is still competitive for single-end data except for compressed outputs where `SeqKit` leads on speed alone.

The 64-bit hashing mode is appropriate for the overwhelming majority of datasets: the collision probability remains below 1‰ for libraries of up to approximately 600 million paired-end reads (1.2 billion total sequences). For larger datasets, the automatic upgrade to 128-bit hashing provides the same pipeline interface with negligible runtime overhead and approximately double the memory cost, a trade-off that remains highly favourable compared to any deterministic alternative.

We advise against `prinseq++` in any production pipeline handling more than 5M reads. Their false-positive rates are not a theoretical concern but an empirically observed failure: they remove correct unique sequences at increasing rates as dataset size grows, silently corrupting downstream variant calling and assembly results. `CD-HIT-DUP`, `FastUniq` and `clumpify` remain valid choices only for small, uncompressed datasets where exact error tolerance or mismatch tolerance are required and where memory is not a constraint. We have to warn about `fastp`, since it use the same bloom filter approach as `prinseq++`, it is likely to suffer from the same false-positive rate explosion at larger volumes, though we did not observe this within the tested range.

### 6.3: Limitations and future work

The current version of `FastDedup` is strictly single-threaded. While this simplifies the zero-allocation pipeline and guarantees deterministic output ordering, it leaves performance on the table for users with access to multiple cores. The most impactful near-term improvement would be parallelizing the compression and decompression stages, which account for more than 90% of wall-clock time in compressed-output modes. Parallelizing the hashing and hash-set lookup stages would offer additional gains, though this requires careful handling of concurrent access to the shared hash set.

A second avenue for development is abundance tracking: rather than discarding duplicate records entirely, `FastDedup` could optionally record the copy number of each unique sequence in the output headers. This would extend its utility to metagenomics and transcriptomics workflows where abundance information carries biological meaning.

We can also note that the current benchmarking suite covers synthetic data derived from a single human reference genome. Extending the evaluation to metagenomic, amplicon, and low-complexity datasets would provide a more complete picture of performance across the full range of sequencing applications. It should be noted that all these benchmarks were performed with `FastDedup` version 1.0.0; improvements and optimizations have been made since then, so the results documented in this paper may not fully reflect the current, likely enhanced performance of `FastDedup`.

Finally, `FastDedup` could continue to expand its feature set to include additional pre-processing steps such as quality filtering, adapter and sequence trimming for example.

`FastDedup` is available under the MIT License at:

- GitHub https://github.com/RaphaelRibes/FastDedup
- Bioconda https://anaconda.org/bioconda/fdedup
- Cargo https://crates.io/crates/fastdedup

The code of the benchmark can be found at: https://github.com/RaphaelRibes/bench-fdedup.git

## 7 Acknowledgements

This paper was written by Raphaël Ribes, reviewed and edited by Céline Mandier, with the assistance of Gemini 3.1 for grammar, and language correction. Code architecture and design was developed by Raphaël Ribes, with autocompletion assisted by Gemini 3.1. Feedback on the algorithm design and ideas for features were provided by Céline Mandier and Alban Mancheron, the implementation was done by Raphaël Ribes. The testsuite was mostly developed by Claude Sonnet 4.6, reviewed by Raphaël Ribes.

Raphaël Ribes was supported through an internship grant awarded to Alice Baniel by the University of Montpellier (I-SITE Excellence Programme 2022 - SOFIT).

Computations were performed on the ISDM-MESO HPC platform, funded in the framework of State-region planning contracts (Contrat de plan État-région – CPER) by the French Government, the Occitanie/Pyrénées-Méditerranée Region, Montpellier Méditerranée Métropole, and the University of Montpellier.

We would like to thank the Data Science Institute of Montpellier (ISDM) funded by the ANR and France 2030 under the UM2030 programme (ANR-21-IDES-0005), for their expertise and their valuable support.

## 8 Annexes

Note – In all Annex tables, values highlighted in green denote tools whose performance is not significantly worse than the best tool in that column. The comparison uses a one-sided Welch’s *t*-test (H0: the tool is no slower than the column-best) with a family-wise error rate of α = 0.05 across all pairwise comparisons within each table. `prinseq++` is excluded from this comparison due to the duplication accuracy issues discussed in Section 5.1 Its values are never highlighted regardless of performance. A tool appearing without green highlighting is significantly slower or more memory-intensive than the column-best at the corrected significance level.

